# Sublethal stress from polypharmacy modulates scavenging function and fenestrations in mouse liver sinusoidal endothelial cells

**DOI:** 10.64898/2026.02.23.707391

**Authors:** Kajangi Gnanachandran, Dina Spigseth Hovland, Jakub Pospíšil, Karolina Szafranska, John Mach, Sarah N. Hilmer, Peter A.G. McCourt

## Abstract

Polypharmacy, the concurrent use of multiple medications, is increasingly prevalent in older people and is associated with adverse outcomes such as falls, frailty, functional and cognitive decline, and increased hospitalization and mortality. The liver, as the primary site of metabolism, is exposed to varying drug concentrations during first pass metabolism, hepatic clearance and perfusion, potentially causing alterations in liver sinusoidal endothelial cells (LSEC). LSEC are specialized endothelial cells responsible for maintaining fenestrations - dynamic, transcellular pores that facilitate the exchange of substances between the blood and liver parenchyma. Disruption of fenestrations can compromise liver function, contributing to a variety of hepatic disorders. This study investigated the effects of four commonly prescribed drugs — metoprolol, citalopram, oxybutynin and oxycodone — on LSEC function. We examined their impact on LSEC viability, endocytosis, and fenestration morphology at both systemic steady-state and first-pass concentrations, separately and in a polypharmacy cocktail to model clinical exposure. All treatments induced sublethal metabolic changes, but effects on LSEC functions were drug- and concentration-dependent. Citalopram and oxybutynin caused dose-dependent defenestration, whereas metoprolol and oxycodone produced mild, non-dose-dependent effects. Endocytic activity was increased with oxybutynin, metoprolol, oxycodone, and the polypharmacy cocktail, while citalopram had no effect. The polypharmacy cocktail triggered synergistic defenestration at first-pass concentrations, but not at steady-state levels. These results highlight the concentration-dependent and combinatorial effects of polypharmacy on LSECs, emphasizing the need to consider endothelial responses in drug safety and pharmacokinetic assessments, particularly in patients exposed to multiple medications.

## 1 Introduction

Polypharmacy, typically defined as the simultaneous use of multiple medications [1], has become an increasingly prevalent phenomenon worldwide due to the ageing global population and the growing burden of chronic diseases [2,3]. It is associated with an elevated risk of adverse drug reactions, drug-drug interactions, functional and cognitive decline and increased hospitalization and mortality [4,5]. The Drug Burden Index (DBI) is a pharmacological risk assessment tool to quantify the functional burden of anticholinergic and sedative medications [6]. Higher DBI scores are often associated with impaired physical function, increased frailty and declines in cognitive function in older adults [7]. A recent study using mouse models showed that chronic exposure to a high-DBI polypharmacy regimen — consisting of citalopram, metoprolol, oxybutynin, oxycodone and simvastatin — increased frailty and impaired physical function [8]. These effects were more severe than those observed with individual drug treatments and were partially reversible through deprescribing (tapered drug cessation) [8].

Among the organs most vulnerable to drug-induced toxicity is the liver, which plays a key role in xenobiotic metabolism and systemic detoxification [9]. During first-pass metabolism, orally administered drugs reach the liver at higher concentrations than those observed systemically, potentially exposing hepatic cells to cytotoxic or sublethal stress [10]. With ageing, the liver’s drug clearance capacity declines due to alterations in hepatic blood flow, metabolic enzymes and hepatic function, compounding the risk of hepatic dysfunction in patients receiving polypharmacy regimens [11,12]. This decrease in hepatic drug metabolism and clearance increases susceptibility to adverse drug reactions and hepatic and systemic drug toxicity in the elderly [13].

Within the liver, liver sinusoidal endothelial cells (LSEC) form a highly specialized barrier between the blood and hepatocytes, characterized by the absence of a basement membrane, high endocytic activity and the presence of fenestrations - transcellular pores that facilitate passive bidirectional transport between the plasma and hepatocytes [14,15]. These fenestrations cover 2-20% of the surface of LSEC [16] and are typically arranged in groups of “sieve plates” delineated by tubulin fibres [17]. Strategically positioned at the front line of hepatic filtration, LSEC are directly exposed to portal blood enriched with drugs absorbed from the gastrointestinal tract [18], and therefore they may encounter higher concentrations of orally administered drugs undergoing first-pass hepatic metabolism [19]. The unique combination of LSEC structural and functional features makes them critical for maintaining hepatic homeostasis, while simultaneously making them particularly sensitive to physiological and pharmacological stressors. Moreover, LSEC fenestrations are supported by an actin cytoskeleton that plays a critical role in maintaining dynamics and responsiveness [20]. Fenestrations can be influenced by various physiological and pharmacological cues, including reactive oxygen species (ROS) [21], calcium signalling and actin-regulatory proteins [22,23], drug exposure and ageing [24]. In ageing and cirrhosis, changes to the fenestrated sinusoidal endothelium may impair transfer of substrates including drugs [12]. While some drugs are known to affect endothelial function [25], the direct, sublethal effects of common medications, especially in the context of polypharmacy, on LSEC fenestrations and function remain poorly characterized. This is especially with respect to sublethal outcomes such as changes in endocytic activity and ultrastructure that may precede cell death.

Here we study young adult mice to reveal the direct impact of chronic polypharmacy independent of ageing, providing insight into early functional and morphological changes that may precede clinical symptoms. Understanding these early effects will contribute towards developing prevention strategies and optimizing medication use that could be validated in future in old age.

In this study, we hypothesized that certain drugs, at sublethal doses, alter LSEC endocytosis and fenestrations, with effects amplified by first-pass concentrations and drug combinations. These alterations may represent early cellular events contributing to hepatic ageing and provide mechanistic insights into how common medications impair liver endothelial function without causing visible cytotoxicity. To test this, we investigated the effects of four commonly prescribed medications — citalopram, oxybutynin, metoprolol and oxycodone — as monotherapies and a high-DBI polypharmacy cocktail, on LSEC function and ultrastructure. We used physiologically relevant concentrations reflecting both systemic steady-state and first-pass exposure, as estimated from *in vivo* mouse studies administering drugs chronically at therapeutic doses [8], and we evaluated their impact on cell viability, endocytic activity and fenestrated morphology.

## 2 Methods

### 2.1 LSEC isolation and cell culture

In this study, all the experiments were carried out in primary LSEC isolated from male C57BL-/-6JRj mice (Janvier Lab, France). The protocols were approved by the local Animal Care and National Animal Research Authority at the Norwegian Food Safety Authority (Mattilsynet; Approval IDS: UiT 20/21 and 09/22). All the procedures were performed in compliance with applicable approved guidelines and regulations and are reported according to the ARRIVE guidelines. Mice were ordered at 5–6 weeks of age and included in experiments after 1–6 weeks of housing under controlled conditions with a standard chow diet (*ad libitum*) with free access to food and water. A minimum of 3 bio replicates were used for each experiment, with a total number of 30 mice.

The mice were sacrificed by cervical dislocation and hepatic cells were extracted by performing a liver perfusion via portal vein as described in detail by Elvevold et al. (2022)[26]. In brief, after the perfusion and digestion with Liberase™ (Sigma-Aldrich, 0.024 mg/mL in isotonic sodium/potassium HEPES buffer at pH 7.4), hepatocytes and non-parenchymal cells (NPCs) were separated by differential centrifugation at 35 × g for 2 min at 4°C, using maximum acceleration and deceleration settings. The resulting supernatant, containing NPCs, was subsequently centrifuged at 300 × g for 8 min at 4°C under the same rotor settings. The NPCs pellet was resuspended in 150 mL of freshly prepared rinsing buffer, MACS BSA Stock Solution diluted 1:20 in autoMACS Rinsing Solution ((Miltenyi Biotec, Germany) and LSEC were isolated from the NPCs fraction using CD146-targeted immunomagnetic separation. Mouse CD146 MicroBeads (Miltenyi Biotec, Germany) were added to the suspension at a 1:20 ratio and incubated for 30 min at 4°C on a rotating mixer. Following this incubation, cells were washed, centrifuged again at 300 × g for 8 min at 4°C, and resuspended in the rinsing buffer. The labelled cell suspension was applied to a pre-washed MS Column (Miltenyi Biotec, Germany) equipped with a 70 µm cell strainer (Miltenyi Biotec, Germany), followed by three washes with 3 mL of rinsing buffer. The column was then removed from the magnetic separator and magnetically retained LSEC were eluted into a fresh collection tube. Isolated cells were then counted and seeded onto wells pre-coated with 0.2 mg/mL human fibronectin.

LSEC were cultured in endothelial cell growth medium (EGM, Cell Applications Inc. USA) supplemented with 10,000 U/mL penicillin (Sigma-Aldrich) and 10 mg/mL streptomycin (Sigma-Aldrich) at 37 °C in 5% CO_2_ and 5% O_2_ overnight before treatment.

### 2.2 Treatments

LSEC were treated with either the steady-state concentration or the first-pass concentration of citalopram (Merck), oxybutynin (Merck), metoprolol (Merck) and oxycodone (Lipomed), both as monotherapies and a high-DBI cocktail containing all four drugs. All drug solutions were prepared as a 10 mM stock solution in DMSO and stored at -80°C for up to 6 months.

After 24h of culture, LSEC were treated with 150 µL of fresh EGM containing the drug solutions at the desired concentration. The samples were kept for 5h at 37°C in 5% CO_2_ and 5% O_2_ before further analysis.

#### 2.2.1 Determination of the steady-state-concentrations

The steady-state concentrations of the drugs used in this study were derived from the *in vivo* experiments conducted by Mach et al. (2021) [8]. Briefly, ageing male C57BL/6 mice received therapeutic doses of drugs; in monotherapy or polypharmacy, in their food/water from 12 to 24 months. The high DBI polypharmacy regimen consisted of simvastatin (20 mg/kg/d), metoprolol (350 mg/kg/d) oxybutynin (27.2 mg/kg/d), oxycodone (5 mg/kg/d) and citalopram (15 mg/kg/d). Identical doses of drug were administered to the monotherapy groups at 24 months of age; blood was collected from submandibular vein and drug levels were measured using liquid chromatography-tandem mass spectrometry method previously described [27]. The steady-state concentrations of monotherapies (Table 1) and polypharmacy cocktails (Table 2) were derived from mean +/- standard error of the mean (unpublished data) derived from measurements at 24 months of age. Although simvastatin was included in the *in vivo* polypharmacy regimen, it was not included in the present *in vitro* study because reliable exposure-matched concentration data for simvastatin were not available, precluding physiologically relevant dose selection.

**Table 1.** steady-state concentration, bioavailability, and estimated first-pass concentration of each drug as monotherapy.

| Drug | Steady-state Concentration (ng/mL) | Bioavailability (%) | First-Pass Concentration (ng/mL) |
| --- | --- | --- | --- |
| Citalopram | 3.54 ± 1.17 | 30 | 11.80 ± 3.90 |
| Metoprolol | 97.23 ± 23.45 | 31 | 313.61 ± 72.55 |
| Oxycodone | 0.15 ± 0.05 | 30 | 0.50 ± 0.17 |
| Oxybutynin | 0.07 ± 0.05 | 57 | 0.12 ± 0.09 |

**Table 2.** steady-state concentration, bioavailability, and estimated first-pass concentration of each drug in the high DBI polypharmacy cocktail.

| Drug | Steady-state Concentration (ng/mL) | Bioavailability (%) | First-Pass Concentration (ng/mL) |
| --- | --- | --- | --- |
| Citalopram | 8.81 ± 1.58 | 30 | 29.37 ± 5.27 |
| Metoprolol | 268.58 ± 47.13 | 31 | 866.71 ± 151.06 |
| Oxycodone | 0.25 ± 0.08 | 30 | 0.83 ± 0.27 |
| Oxybutynin | 0.13 ± 0.06 | 57 | 0.23 ± 0.11 |

#### 2.2.2 Calculation of the first-pass concentrations

The first-pass concentrations of each drug were calculated using the ratio of its steady-state concentration to its oral bioavailability [28]. The bioavailability values in rodents for citalopram were obtained from Overø (1982) [29], for metoprolol from Yoon et al. (2011) [30] and for oxycodone from Raleigh et al. (2018) [31]. The bioavailability of oxybutynin in humans is approximately 6%, with an increase to 31% when taken with food [32,33]. There is no published data on the bioavailability of oxybutynin in rats or mice, and the correlation of bioavailability of drugs in rodents compared to humans is not strong [34]. Therefore, the higher end of the human bioavailability (31%) was used as a reference for this drug.

The table below contains the summary of the concentrations for the specific drugs used in this study.

In case of the high DBI polypharmacy cocktail, the concentrations were almost double those obtained after the monotherapies. The table below lists these values.

### 2.3 Viability assay - Resazurin

Cells were cultured on standard 48-well plates (60,000 cells/well) and their mitochondrial function and viability were assessed using a commercially available Resazurin/resorufin assay (Biotechne). Cells were first pre-treated with the desired concentrations of the drugs for 2 h, and then the volume was reduced to 90 µL per well and 1:10 resazurin reagent was added to the culture media for 3h, making a total of 5 h treatment time. After this timepoint, 50 µL of supernatant was collected and the fluorescent signal was measured using a plate reader (ClarioStar, BMG Labtech) with the excitation set at 530–570 nm and emission at 580–590 nm.

### 2.4 Scavenging assay

For quantitative studies of endocytosis and degradation in LSEC, a radiolabelled formaldehyde-treated serum albumin (^125^I-FSA)-based scavenging assay was used. Cells were cultured on standard 48-well plates at a high density (200.000 cells/well) and, after a pre-treatment of 2 h with the desired concentrations of the drugs, around 30 ng of ^125^I-FSA was added to each well together with 1% human serum albumin (HSA) (Alburex, CSL Behring) and incubated for 3 h. Subsequently, the radioactivity of the cell-associated and degraded FSA fractions was measured using a gamma counter (Packard Cobra Gamma, GMI Inc) and analysed according to previously described methods [35,36].

### 2.5 Scanning electron microscopy

For imaging with scanning electron microscopy (SEM), cells were cultured on fibronectin-coated 16-well plates (CS16-CultureWell™ Removable Chambered 126 Coverglass, Grace Bio-labs) with a density of 60.000 cells/well. After 5 h of treatment, they were fixed with 4% formaldehyde and 1% glutaraldehyde in PHEM buffer comprised of 60 mM PIPES (Sigma-Aldrich), 25 mM HEPES (Thermo Fisher), 10 mM EDTA (Sigma-Aldrich) and 2 mM MgCl_2_ (Merck), pH 7) and stored at 4°C until further processing. The fixed samples were washed 3 times with PHEM buffer and incubated with freshly made 1% tannic acid (Sigma Aldrich) in PHEM buffer for 1 h. Samples were washed again three times with PHEM buffer, then incubated with 1% osmium tetroxide (Electron Microscopy Sciences) in ddH_2_O for 30 min, followed by another triple wash with PHEM. Dehydration was performed using a graded ethanol series: 30%, 60%, and 90% for 5 min each, followed by 100% ethanol for 4 × 5 min. A chemical drying with hexamethyldisilazane (Sigma-Aldrich) was applied twice for 2 min, and samples were dried overnight in a desiccator. The processed samples were mounted on aluminium stubs and glued on using a silver glue (Agar Scientific). Glue was allowed to dry for at least 3 h. Just before imaging, samples were sputter-coated with a 10 nm layer of palladium/gold alloy (Quorum). The images were taken using the Zeiss Gemini 300 scanning electron microscope. For each biological replicate, images were taken from 15–20 cells in three independent and randomly chosen areas, with a total of 45–60 images/experimental group.

### 2.6 Quantitative image analysis

A custom StarDist [37,38] segmentation model using star-convex polygon representations was trained to identify fenestrations in LSEC from scanning electron microscopy images. The U-Net-based model [39] was trained on 22 manually annotated images and validated on 11 images, enabling it to accurately identify fenestrations. This segmentation enabled quantitative analysis of fenestration characteristics including fenestration frequency (number of fenestrations per cell area µm^2^), porosity (% of fenestration-occupied cell area) and diameter of the fenestration present in each cell. Our implementation utilized the publicly available StarDist GitHub repository (https://github.com/stardist/stardist) with methodology adapted from the original framework [37].

### 2.7 Statistics

All data is presented as mean ± standard deviation (STD). All values from treated samples were normalized to the corresponding untreated control (0.01% DMSO), which was set to 100%. Normality of distribution was assessed using Shapiro-Wilk test. Data with normal distribution was analysed by One-way ANOVA followed by Dunnet’s multiple comparisons test. Data with abnormal distribution was analysed by Kruskal-Wallis test followed by Dunn’s multiple comparisons test. All analyses were performed using GraphPad Prism version 10.2.4 for Windows (Dotmatics, GraphPad Software, San Diego, CA, USA), where a significance level of *P* −*values* = 0.05 was accepted as significant. Significance levels were denoted using the conventional asterisk annotation: *p* ≥ 0.05 = ns, *p <* 0.05 = *, *p <* 0.01 = **, *p <* 0.001 = ***. Detailed statistical results are provided in Supplementary Tables S1-S3.

## 3 Results

### 3.1 First-pass concentrations of drugs enhance LSEC scavenging function without affecting their viability

To assess potential cytotoxic effects of mono- and polypharmacy treatments on liver sinusoidal endothelial cells (LSEC), cell viability was evaluated using the resazurin assay, which reflects cellular metabolic activity and reduction potential (Figure 1A–B). Treatment with steady-state concentrations of individual drugs (citalopram, oxybutynin, metoprolol, and oxycodone) or the high DBI polypharmacy cocktail did not significantly alter viability, with values ranging from 97% to 106% of control (*p >* 0.05, Figure 1A). Similarly, first-pass concentrations of the monotherapies caused a slight, but non-significant, increase in viability readouts (109%–122% of control, *p >* 0.05, Figure 1B). Furthermore, the cells treated with first-pass concentrations of the polypharmacy cocktail processed resazurin significantly more than the untreated samples (128%, *p <* 0.05, Figure 1B). These results indicate that none of the treatments compromised LSEC viability regardless of the concentration used, except for the higher concentration of the polypharmacy cocktail.

**Fig. 1.**
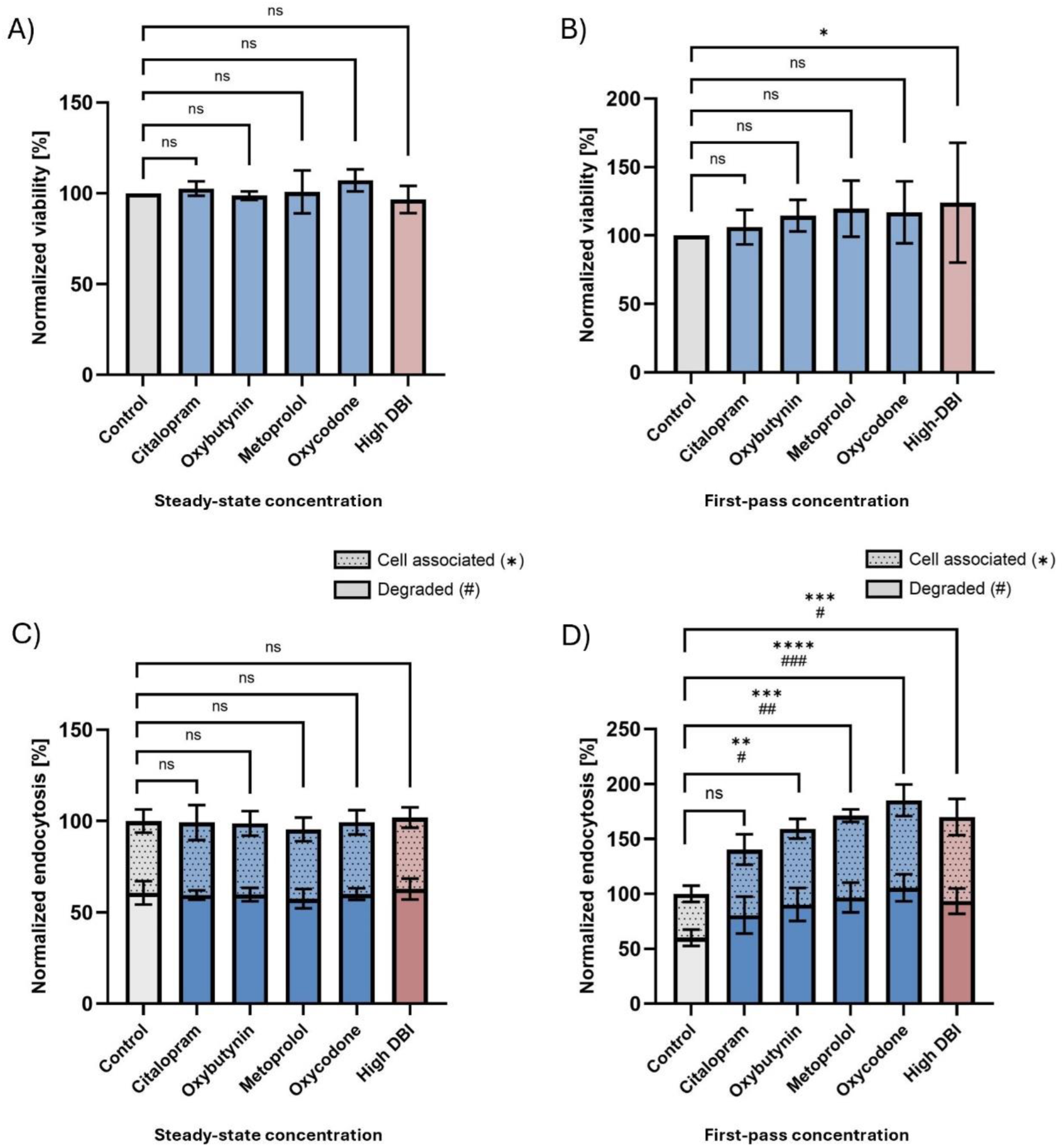
Effects of polypharmacy on LSEC viability and scavenging function. (A,B) LSEC viability after 5 h exposure to citalopram (3.54 and 11.80 ng/mL), oxybutynin (0.07 and 0.12 ng/mL), metoprolol (97.23 and 313.61 ng/mL), oxycodone (0.15 and 0.50 ng/mL), or the high DBI polypharmacy cocktail at steady-state (concentrations were approximately two-fold higher than those observed during monotherapy) (A) or first-pass (B) concentrations. (C,D) Scavenger receptor-mediated endocytosis of ^125^I-FSA measured after exposure to the same drug concentrations at steady-state (C) or first-pass (D). Light bars indicate cell-associated ligand and dark bars indicate degraded ligand. Data are normalized to the untreated control (0.01% DMSO) and are expressed as mean ± standard deviation of 3-4 independent biological replicates. Statistical significance is indicated in the graphs (*p <* 0.05 = *, *p <* 0.01 = **, *p <* 0.001 = ***)

To evaluate potential functional effects on LSEC, endocytic activity was measured using ^125^I-labelled formaldehyde-treated serum albumin (^125^I-FSA), which is degraded after internalization_by scavenger receptors (Figure 1C–D). At steady-state concentrations, neither individual drugs nor the polypharmacy cocktail affected total FSA uptake or degradation, with cell-associated and degraded fractions remaining comparable to control levels (Figure 1C). However, all treatment with first-pass concentrations, except citalopram, significantly enhanced LSEC endocytic function (Figure 1D). Total endocytosis increased to 160% with oxybutynin (*p <* 0.01), 171% with metoprolol (*p <* 0.001), and 185% with oxycodone (*p <* 0.0001), compared to solvent control. The polypharmacy cocktail also significantly increased total uptake to 170% of control (*p <* 0.001).

All the normalized values, with the respective statistical analysis, are given in table S1 and table S2 of the supplementary material. All data represents mean ± standard deviation (STD) from at least three independent experiments. These data suggest that although cell viability remains unaffected, first-pass drug exposure enhances the scavenging function of LSEC in all treatments except for citalopram, possibly reflecting an early functional response to sublethal stress.

### 3.2 Effects of mono- and polypharmacy treatments on LSEC morphology

To investigate the effect of drug exposure on the ultrastructural organization of LSEC fenestrations, we used SEM alongside quantitative morphometric analyses. Fenestration architecture was characterized using two principal metrics: fenestration frequency, defined as the number of fenestrations per unit surface area, and porosity, denoting the proportion of the cellular surface occupied by fenestrations. Under physiological conditions, LSEC are characterized by fenestrations that cover 2-20% of their surface, typically arranged in groups as “sieve plates” [16]. Upon pharmacological treatments, we observed a continuum of morphological alterations affecting these parameters.

#### 3.2.1 Impact of steady-state-level drug exposure on LSEC porosity and fenestration frequency

Figure 2A shows representative SEM images of LSEC treated with steady-state concentrations of citalopram, oxybutynin, metoprolol, oxycodone, and the high DBI polypharmacy cocktail. Visual inspection suggests that all monotherapies and the polypharmacy treatment preserved overall fenestration patterns, with no dramatic loss of pores compared to the solvent control. However, subtle differences in fenestration organization, such as sieve plate grouping, were noted, which are more clearly captured by the quantitative analysis described below.

**Fig. 2.**
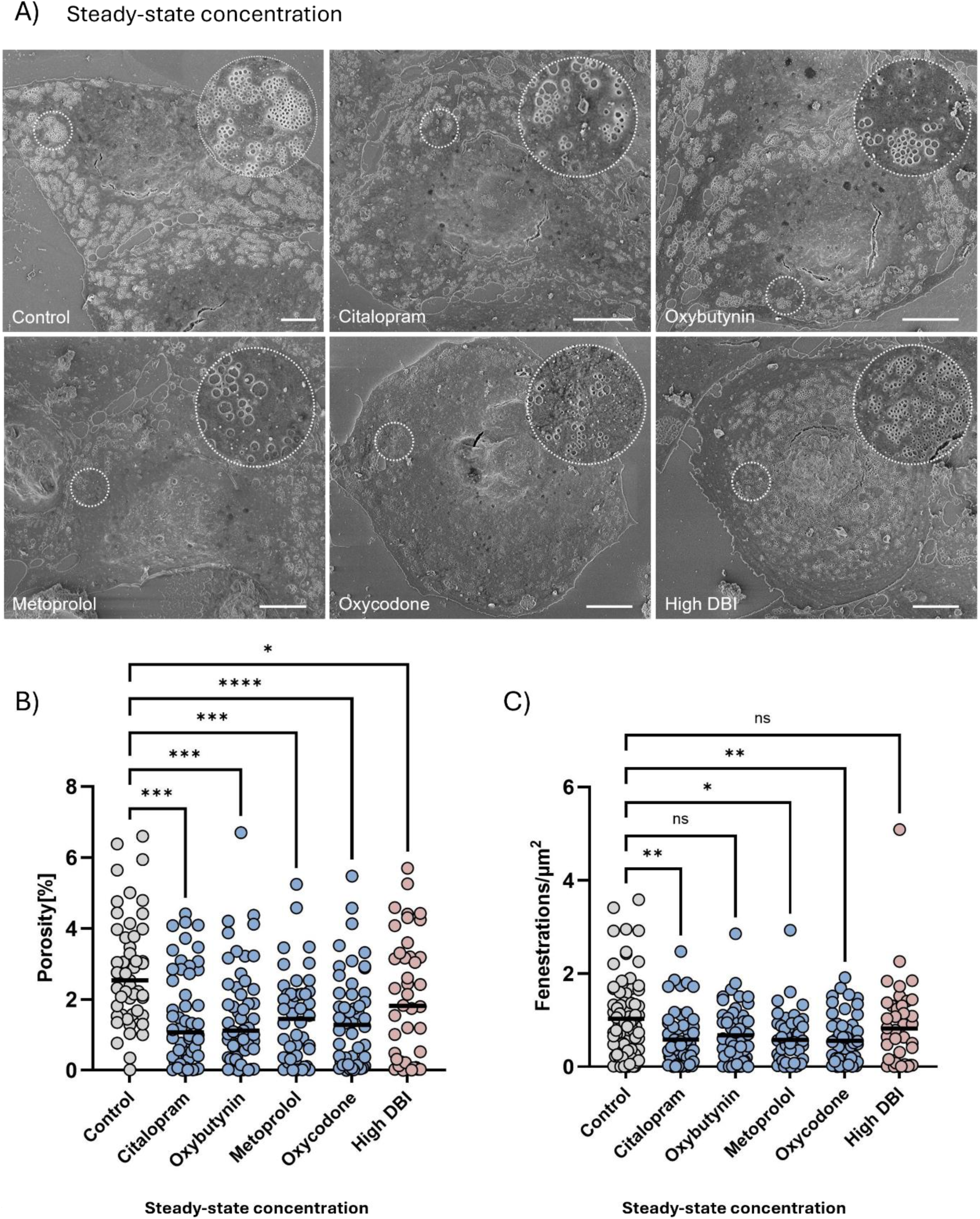
Effects of steady-state concentration polypharmacy on LSEC fenestrated morphology. (A) Representative scanning electron microscopy images of LSEC after 5 h treatment with citalopram (3.54 and 11.80 ng/mL), oxybutynin (0.07 and 0.12 ng/mL), metoprolol (97.23 and 313.61 ng/mL), oxycodone (0.15 and 0.50 ng/mL), or the high DBI polypharmacy cocktail at steady-state (concentrations were approximately two-fold higher than those observed during monotherapy). Scalebar = 5 µm and inset diameter = 4 µm. (B,C) Quantitative analysis of fenestrated morphology: porosity (percentage of cell surface covered by fenestrations, B) and fenestration frequency (number of fenestrations per µm^2^, C). Each dot represents a single cell (45–60 per treatment), and black lines indicate mean values from 3–4 independent biological replicates. Data is normalized to the untreated control (0.01% DMSO). Statistical significance was assessed using Kruskal-Wallis test with Dunn’s multiple comparisons; significance levels are denoted using the conventional asterisks annotation: *p* ≥ 0.05 = ns, *p <* 0.05 = *, *p <* 0.01 = **, *p <* 0.001 = ***.

Quantitative analysis of porosity (percentage of the cell area covered with fenestrations, Figure 2B) revealed that all the treatments significantly reduced LSEC porosity compared to the solvent control (2.31 ± 1.54). Specifically, citalopram with a mean porosity of 1.49 ± 1.34 (*p <* 0.001), oxybutynin with 1.57 ± 1.48 (*p <* 0.001), metoprolol with 1.42 ± 1.26 (*p <* 0.001), and oxycodone with 1.42 ± 1.35 (*p <* 0.0001). The polypharmacy cocktail, while still statistically significant, exhibited the least reduction in porosity with a mean of 2.01 ± 1.74 (*p <* 0.05). Fenestration frequency (number of fenestrations per µm^2^, Figure 2C) followed a similar trend. Relative to the control (1.03 ± 0.81), the most pronounced decreases were observed with citalopram and oxycodone, both yielding mean frequencies of 0.58 ± 0.60 (*p <* 0.01) and 0.58 ± 0.53 (*p <* 0.01) respectively. Metoprolol also reduced fenestration frequency (0.58 ± 0.54, *p <* 0.05), whereas oxybutynin and the polypharmacy cocktail did not produce significant changes at steady-state concentration.

Collectively, these findings indicate that, at steady-state concentrations, certain drugs substantially reduce LSEC fenestration frequency and porosity, while others exert minimal effects. Interestingly, in the polypharmacy regimen, fenestrations appear less affected than would be expected from the effects of the individual drugs, suggesting potential drug-drug interactions that modulate endothelial responses; this phenomenon is most apparent at steady-state rather than during first-pass exposure and is discussed further in the next section.

#### 3.2.2 Effects of first-pass concentrations on LSEC porosity and fenestration frequency

To assess the impact of higher, pre-hepatic drug concentrations, LSEC were also exposed to first-pass levels of the same treatments. The SEM images represented in Figure 3A, in contrast to steady-state-level exposures, show visibly reduced fenestrations across all treatment groups, suggesting a more pronounced disruption of fenestration architecture.

**Fig. 3.**
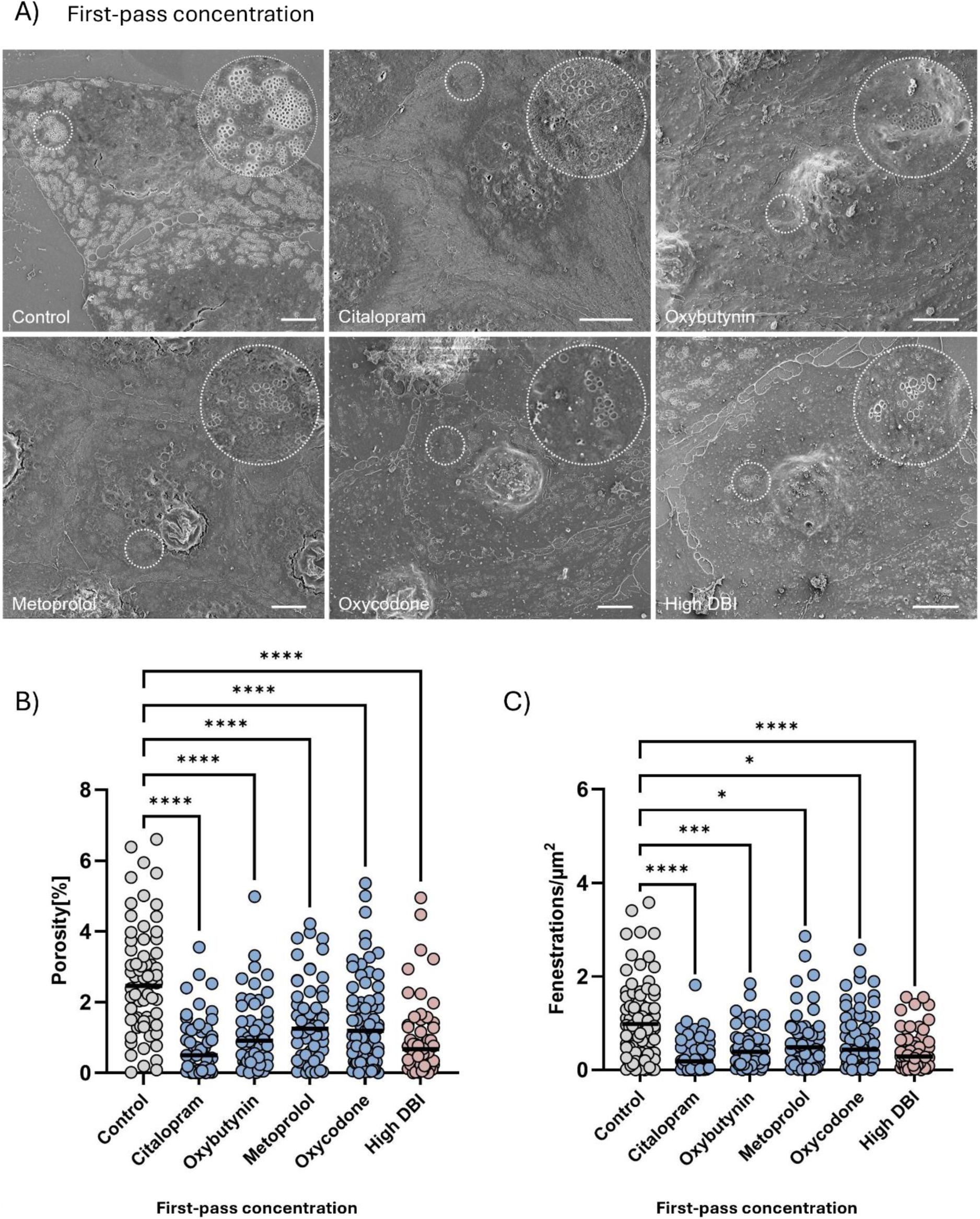
Effects of first-pass concentration polypharmacy on LSEC fenestrated morphology. (A) Representative scanning electron microscopy images of LSEC after 5 h treatment with citalopram (3.54 and 11.80 ng/mL), oxybutynin (0.07 and 0.12 ng/mL), metoprolol (97.23 and 313.61 ng/mL), oxycodone (0.15 and 0.50 ng/mL), or the high DBI polypharmacy cocktail at steady-state (concentrations were approximately two-fold higher than those observed during monotherapy). Scalebar = 5 µm and inset diameter = 4 µm. (B,C) Quantitative analysis of fenestrated morphology: porosity (percentage of cell surface covered by fenestrations, B) and fenestration frequency (number of fenestrations per µm^2^, C). Each dot represents a single cell (45–60 per treatment), and black lines indicate mean values from 3–4 independent biological replicates. Data is normalized to the untreated control (0.01% DMSO). Statistical significance was assessed using Kruskal-Wallis test with Dunn’s multiple comparisons; significance levels are denoted using the conventional asterisks annotation: *p* ≥ 0.05 = ns, *p <* 0.05 = *, *p <* 0.01 = **, *p <* 0.001 = ***.

Quantitative analysis confirmed these observations. Porosity (Figure 3B) was significantly reduced in all treatment groups compared to the solvent control (2.61 ± 1.54), with mean values of 0.75 ± 0.83 for citalopram, 1.09 ± 1.0 for oxybutynin, 1.42 ± 1.18 for metoprolol, 1.44 ± 1.27 for oxycodone, and 0.97 ± 1.10 for the polypharmacy cocktail, all with *p <* 0.0001. Fenestration frequency (Figure 3C) was also markedly reduced compared to the solvent control (0.99 ± 0.79). Mean frequency values were 0.31±0.37 for citalopram (*p <* 0.0001), 0.48± 0.43 for oxybutynin (*p <* 0.001), 0.63 ± 0.62 for metoprolol (*p <* 0.05), 0.63 ± 0.59 for oxycodone (*p <* 0.05), and 0.42 ± 0.43 for the polypharmacy cocktail (*p <* 0.0001).

These results demonstrate that first-pass drug concentrations induce a consistent and significant reduction in fenestration frequency and porosity in LSEC, highlighting the potentially greater impact of elevated local drug levels encountered during hepatic first-pass exposure.

To further characterize the impact of drug exposure on LSEC morphology, fenestration diameter was quantified under both steady-state-level and first-pass treatment conditions. The results are shown in Figure 4.

**Fig. 4.**
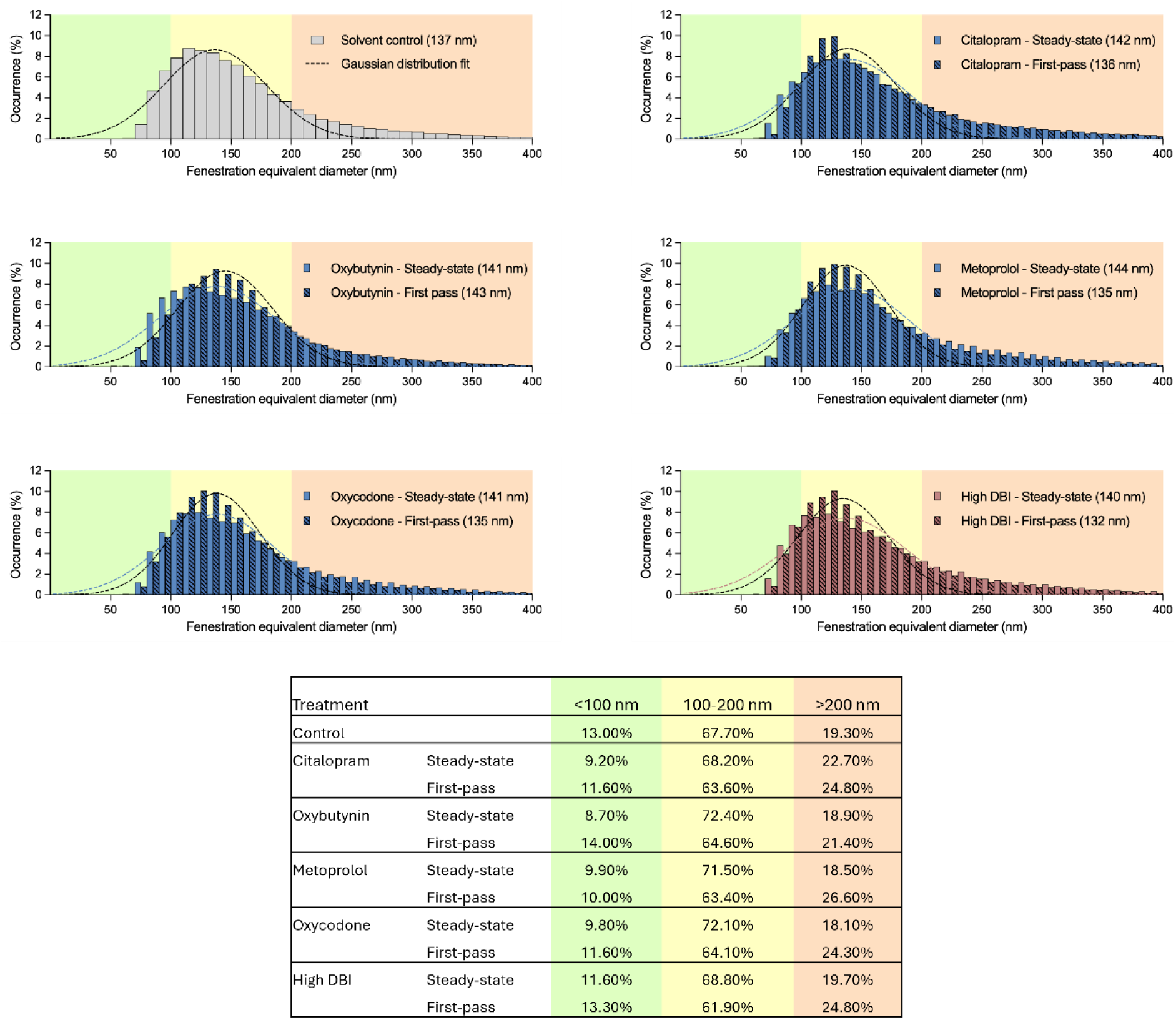
Distribution of fenestration diameter of LSEC treated with steady-state and first-pass concentrations of drugs. Each histogram represents equivalent diameter measurements of fenestrations from at least 3 biological replicates. The dashed lines represent the fitted Gaussian curves. The centre of each Gaussian distribution is noted in the legend for each treatment group separately. The table below summarizes the percentage of total fenestrations per condition within three predefined diameter ranges: small (<100 nm, green), intermediate (100-200 nm, yellow) and large (>200 nm, orange). The single distribution of fenestration diameter for each treatment group is presented in the supplementary figures S1 and S2.

Fenestration diameter analysis revealed no significant drug-specific patterns. However, at first-pass concentrations, all treatments exhibited an increased fraction of large fenestrations (>200 nm) compared to the control, indicative of early structural alterations. In contrast, at steady-state concentrations, the proportion of small fenestrations (<100 nm) was consistently reduced across all treatments, while intermediate fenestrations (100—200 nm) remained largely unchanged.

## 4 Discussion and conclusions

Polypharmacy is increasingly recognized as a contributor to adverse drug reactions, frailty and functional decline, particularly in ageing populations [1,5]. Despite the central role of the liver in drug metabolism, the direct sublethal effects of common medications on LSEC remain poorly characterized [40]. This study aimed to investigate whether physiologically relevant drug exposures, both at steady-state and first-pass concentrations, alter LSEC viability, endocytosis or fenestration morphology, potentially contributing to early hepatic dysfunction. Our study revealed distinct patterns of LSEC response across treatments and concentrations. While steady-state-level exposures induced moderate changes, first-pass concentrations consistently triggered more pronounced alterations, in particular in endocytosis and fenestration morphology. These results are discussed in detail below.

### 4.1 Viability

LSEC viability was assessed using a resazurin assay, which measures mitochondrial metabolic activity by quantifying the reduction of resazurin to the fluorescent compound resorufin. Across all treatment groups, no significant cytotoxic effects were observed at either steady-state or first-pass concentrations (Figure 5), indicating that the drugs have sublethal effects on LSEC.

Interestingly, first-pass exposure to the high-DBI polypharmacy cocktail resulted in a marked increase in resazurin reduction, suggesting enhanced mitochondrial activity rather than true increase in viability. This apparent increase may reflect a stress-induced metabolic shift rather than improved cell health. All four drugs in the cocktail have been independently associated with oxidative stress and excess reactive oxygen species (ROS) production in various cellular models [41–44]. A similar increase in reducing potential detected with resazurin was observed in LSEC treated with low concentration of hydrogen peroxide (0.5 µM) [21]. It has been proposed that polypharmacy may lead to mitochondrial dysfunction in a range of organs [45]. Moreover, mitochondrial ROS can elevate intracellular calcium levels, which in turn activate calcium-dependent dehydrogenases involved in ATP synthesis [46,47]. These dehydrogenases stimulate the electron transport chain, enhancing mitochondrial respiration and increasing cellular energy output, thereby accelerating resazurin reduction [48,49]. Although these studies were conducted in cardiac cells, similar mechanisms can possibly occur in liver sinusoidal endothelial cells, providing a potential explanation for the observed acceleration of resazurin reduction in our experiments. Thus, the elevated fluorescence signal may not indicate improved viability, but rather a drug-induced metabolic response, potentially driven by ROS and calcium signalling.

These findings are particularly relevant in the context of ageing, where increased mitochondrial ROS and impaired calcium homeostasis contribute to endothelial dysfunction and reduced hepatic resilience [50]. The *in vivo* model of polypharmacy resulted in changes to the hepatic proteome in mitochondrial, antioxidant, inflammatory and immune pathways [51]. Our results suggest that polypharmacy may mimic ageing-like stress responses in LSEC, even in young adult mouse models, highlighting a potential mechanism for premature endothelial decline.

### 4.2 Endocytosis

To assess LSEC endocytic function, we used an iodine-labelled formaldehyde-treated serum albumin (^125^I-FSA) assay. This method quantifies both cellular uptake and degradation of FSA, a ligand internalized primarily via the endocytic receptors stabilin-1 and -2 expressed on LSEC [52].

Consistent with the viability assay, drug treatments at steady-state-level concentrations did not significantly affect endocytic activity, suggesting that systemic exposure is insufficient to disrupt stabilin-mediated uptake. First-pass concentrations of oxybutynin, metoprolol, oxycodone and the polypharmacy cocktail significantly enhanced endocytosis. In contrast, citalopram did not increase endocytosis at either concentration, highlighting a unique functional response among the tested drugs. This pattern suggests drug-specific modulation of LSEC function, potentially linked to intracellular stress signalling.

One plausible mechanism involves ROS, which, as mentioned earlier, are known to elevate intracellular calcium levels. Increased calcium can activate calcium/calmodulin-dependent protein kinase kinase beta (CaMKK-*β*), which in turn signals through protein kinase C zeta (PKC-*ζ*) and AMP-activated protein kinase (AMPK) to enhance clathrin-mediated endocytosis in various cell types [16,53,54]. Although this pathway has not been fully characterized in LSEC, CaMKK-*β*, PKC-*ζ* and AMPK are expressed in these cells and contribute to metabolic regulation and cytoskeletal remodelling [16,55]. Given that stabilin-1/-2-mediated FSA uptake occurs via clathrin-mediated endocytosis [56,57], it is plausible that drug-induced oxidative stress triggers this calcium-dependent pathway, thereby enhancing endocytic activity. Unlike the other drugs, citalopram has a selective serotonergic activity, and it increases extracellular serotonin levels [41], which may selectively remodel fenestrations through cytoskeletal pathways without sufficiently activating the Ca^2+^-dependent CME pathway.

Interestingly, the polypharmacy cocktail did not produce a more pronounced increase in scavenging than the individual drug monotherapies. This may indicate that each monotherapy already reaches a threshold sufficient to maximally stimulate endocytosis, leaving limited capacity for further enhancement. Alternatively, pharmacodynamic interactions among the drugs may dampen or counterbalance each other’s effects, preventing additive responses. Moreover, the drug-induced increase in endocytosis observed here may reflect a compensatory response to sublethal stress, mimicking early ageing-like adaptations. Such hyperactivation may temporarily preserve scavenger function, but sustained stimulation could lead to lysosomal overload, receptor dysregulation and eventual exhaustion of the endocytic machinery.

### 4.3 Morphology

The effects of pharmaceutical treatments on LSEC morphology were assessed using scanning electron microscopy (SEM), enabling direct visualization of fenestrations. Quantitative analysis was performed using a custom StarDist-based segmentation pipeline to extract porosity (percentage of cell surface occupied by fenestrations), fenestration frequency (fenestration per µm^2^) and diameter (pore size distribution).

At steady-state concentrations, all the monotherapies caused reductions in porosity, while fenestration frequency was generally maintained. These subtle changes corresponded to shifts in fenestration size distribution: small fenestration (<100 nm) decreased slightly, intermediate fenestrations (100-200 nm) remained largely unchanged or slightly increased. This indicates early redistribution of fenestration sizes rather than widespread loss of pores at steady-state levels. These patterns are in line with previously reported effects of caffeine, where shifts in fenestration size reflected early endothelial responses to drug exposure [58]. Interestingly, only the high DBI cocktail at steady-state concentration preserved normal fenestration architecture. In contrast, first-pass drug concentrations produced more profound morphological changes. All the treatments significantly reduced porosity and, in most cases, also fenestration frequency, consistent with stronger disruption of sieve plates. Specifically, citalopram, oxybutynin and the polypharmacy cocktail caused marked defenestration, reducing both porosity and fenestration frequency while increasing the proportion of large fenestrations, suggesting that disappearance of sieve plates was driven primarily by loss of intermediate pores. Metoprolol and oxycodone preserved fenestration frequency and showed no clear concentration-dependent effect. This defenestration could reduce transfer of drugs across the hepatic sinusoid to the hepatocytes, reducing first pass hepatic clearance, as demonstrated for diazepam clearance *in vivo* in old age and with Poloxymer 407-mediated defenestration [59].

It is worth noting that fenestration loss was more pronounced at the higher concentrations for citalopram and oxybutynin, but not for metoprolol or oxycodone (Figure 4C). While mechanistic conclusions cannot be drawn from two concentrations alone, the differential concentration sensitivity observed for citalopram and oxybutynin raises the possibility of distinct structural remodelling responses compared to metoprolol and oxycodone. Importantly, all four drugs have been reported to induce ROS generation, albeit to varying degrees and through distinct mechanisms [41–44]. Previous work demonstrated that hydrogen peroxide induced ROS formation can hamper fenestration dynamics and eventually lead to complete defenestration and cell death [21,60]. The lack of reduction in endocytic activity, however, suggest that the tested drugs affect LSEC also via other molecular mechanisms. This shared property provides a unifying mechanistic framework for interpreting the observed morphological changes as follows.

Among the drugs tested in this study, metoprolol and oxycodone caused mild, non–concentration-related fenestration closure, whereas citalopram uniquely induced a uniform shrinkage. At higher concentrations, drug combinations triggered synergistic defenestration. Notably, citalopram dissociated morphological remodelling from functional uptake, indicating that structural and functional alterations in LSEC are not necessarily coupled. This distinction highlights that pharmacological stress can differentially affect structural and functional LSEC endpoints, underscoring the importance of considering both dimensions when interpreting endothelial responses. Ca^2+^ is known to play a central role in regulating LSEC fenestrations through pathways involving myosin light chain kinase, protein kinase C, and RhoA–ROCK signalling [22], while reactive oxygen species (ROS) can exacerbate defenestration by perturbing cytoskeletal integrity under conditions of oxidative stress [16]. Elucidating the spatial and temporal dynamics of these signals in drug-challenged LSEC would require their simultaneous assessment alongside fenestration architecture. This would require advanced super-resolution optical imaging approaches [61] although live cell imaging of LSEC fenestrations with these techniques remains challenging due to ROS induced phototoxicity [60].

## 5 Conclusions

This study demonstrates that commonly used drugs, such as citalopram, oxybutynin, metoprolol and oxycodone, differentially modulate LSEC structure and function depending on the exposure level. All drugs likely induced sublethal changes, but their functional and morphological effects were distinct. Citalopram and oxybutynin caused more profound morphological changes at first-pass concentrations compared to steady-state levels, whereas metoprolol and oxycodone produced only mild fenestration loss with little difference between the two concentrations. In contrast, endocytic activity was enhanced by all drugs at higher concentrations, modelling exposure during first pass metabolism, while citalopram had no significant effect on endocytosis despite inducing defenestration at first-pass concentration. This uncoupling of structural and functional responses suggests that serotonin-mediated cytoskeletal remodelling may alter pore architecture without engaging endocytic signalling pathways.

The effects of the polypharmacy cocktail highlighted the importance of exposure level: at steady-state concentrations, it did not significantly affect morphology, whereas first-pass concentrations triggered synergistic defenestration, exceeding the LSEC capacity to maintain normal fenestration architecture. Drugs that increase endocytosis without fenestration loss may maintain LSEC clearance function, whereas drugs or combinations that cause fenestration loss, such as citalopram, oxybutynin or the high DBI polypharmacy cocktail, could impair microcirculation and macromolecular clearance. Importantly, these *in vitro* findings may provide a mechanism for the *in vivo* study by Mach et al. [8], in which chronic polypharmacy in aged mice exacerbated frailty and impaired physiological function. While our *in vitro* model specifically captures direct LSEC responses to drug exposure, the *in vivo* data suggest that endothelial alterations of this type may represent one of several contributing factors to the systemic effects associated with polypharmacy, including impaired liver function and altered drug metabolism [51].

These results emphasize the physiological relevance of sublethal endothelial responses in polypharmacy, the concentration-dependent nature of drug effects and the importance of considering hepatic endothelial health in drug safety and pharmacokinetic assessments. Future studies should validate these findings in females and cells derived from old animals and investigate the underlying molecular mechanism to better predict and mitigate drug-induced hepatic consequences in patients receiving multiple medications.

## Author Contributions

KG: Conceptualization, Methodology, Investigation, Formal Analysis, Visualization, Writing - Original Draft

DSH: Methodology, Investigation, Formal Analysis, Visualization, Writing - Review and Editing

JP: Quantitative analysis of SEM images, Visualization - Writing - Review and Editing

KS: Methodology, Technical support, Writing - Review and Editing

JM-SH: Determination of the drug concentrations, Writing - Review and Editing

PM: Funding acquisition, Conceptualization, Writing - Reviewing and Editing

## Acknowledgments

This study was supported by EU project DeLIVERy EIC-2021-Pathfinder grant no. 101046928. J.M. was funded by the Penney Ageing Research Unit, Royal North Shore Hospital, Australia.

