## Supplementary data for "Sublethal stress from polypharmacy modulates scavenging function and fenestrations in mouse liver sinusoidal endothelial cells"

***Table S1**: results and statistical significance for Figure 1A and 1B. Using Shapiro-Wilk's test, the distribution of the results was determined to be abnormally distributed for serum concentrations and normally distributed for first pass concentrations. Therefore, Kruskal Wallis test with Dunn's multiple comparisons test was used to identify the statistical significance for serum concentration, and One-Way Anova with Dunnet's multiple comparisons test was used for first pass concentrations.*

| | | Viability (% of control) | | | |
|---|---|---|---|---|---|
| | | *Mean* | *STD* | *P-value* | * |
| **[SERUM]** | **Control** | 100 | | | |
| | **Citalopram** | 101.8 | 14.25 | >0.9999 | ns |
| | **Oxybutynin** | 99.5 | 14 | >0.9999 | ns |
| | **Metoprolol** | 100.9 | 15.51 | >0.9999 | ns |
| | **Oxycodone** | 106.4 | 15.03 | 0.7203 | ns |
| | **High DBI** | 96.73 | 17.95 | >0.9999 | ns |
| **[FIRST PASS]** | **Control** | 100 | | | |
| | **Citalopram** | 109.3 | 35 | 0.7707 | ns |
| | **Oxybutynin** | 116.1 | 25.75 | 0.2888 | ns |
| | **Metoprolol** | 121.5 | 30.31 | 0.0865 | ns |
| | **Oxycodone** | 119.4 | 33.18 | 0.1439 | ns |
| | **High DBI** | 128.4 | 50.99 | 0.0128 | * |

***Table S2:*** *results and statistical significance for Figure 1C and 1D. Using Shapiro-Wilk's test, the distribution of the results was determined to be abnormally distributed for cell associated $I^{125}$-FSA for serum concentrations and normally distributed for degraded ligand after serum treatment, and for both cell associated and degraded ligand after first pass treatments. Therefore, Kruskal Wallis test with Dunn's multiple comparisons test was used to identify the statistical significance for CA serum concentration, and One-Way Anova with Dunnet's multiple comparisons test was used for the remaining parameters.*

| | | Endocytosis (%of control) | | | | | | | |
|---|---|---|---|---|---|---|---|---|---|
| | | ***Cell associated*** | | | | ***Degraded*** | | | |
| | | *Mean* | *STD* | *P-value* | * | *Mean* | *STD* | *P-value* | * |
| **[SERUM]** | **Control** | 39.31 | 6.39 | | | 60.69 | 6.39 | | |
| | **Citalopram** | 39.71 | 9.54 | >0.9999 | ns | 59.49 | 2.56 | >0.9999 | ns |
| | **Oxybutynin** | 39.06 | 6.7 | >0.9999 | ns | 59.72 | 3.67 | 0.9936 | ns |
| | **Metoprolol** | 37.93 | 6.51 | >0.9999 | ns | 57.53 | 5.32 | >0.9999 | ns |
| | **Oxycodone** | 39.34 | 6.65 | >0.9999 | ns | 60.02 | 3.13 | 0.9973 | ns |
| | **High DBI** | 39.08 | 5.58 | >0.9999 | ns | 62.81 | 5.74 | 0.2267 | ns |
| **[FIRST PASS]** | **Control** | 40.06 | 7.43 | | | 59.94 | 7.43 | | |
| | **Citalopram** | 59.7 | 13.87 | 0.0882 | ns | 80.57 | 16.83 | 0.2199 | ns |
| | **Oxybutynin** | 69.88 | 9 | 0.0045 | ** | 90.29 | 15.04 | 0.0206 | * |
| | **Metoprolol** | 74.52 | 5.68 | 0.0006 | *** | 96.64 | 13.53 | 0.0036 | ** |
| | **Oxycodone** | 79.47 | 14.4 | <0.0001 | **** | 105.55 | 12.26 | 0.0003 | *** |
| | **High DBI** | 76.58 | 16.49 | 0.0002 | *** | 93.36 | 11.47 | 0.0129 | * |

***Table S3:*** *results and statistical significance for Figure 2 and 3. Using Shapiro-Wilk's test, the distribution of the results was determined to be abnormally distributed for all of the data. Therefore, Kruskal Wallis test with Dunn's multiple comparisons test was used to identify the statistical significance for both porosity and fenestration frequency of both concentrations.*

| | | Morphology | | | | | | | |
|---|---|---|---|---|---|---|---|---|---|
| | | ***Porosity (% fenestration covered area)*** | | | | ***Fenestration frequency (fen/µm^2)*** | | | |
| | | *Mean* | *STD* | *P-value* | * | *Mean* | *STD* | *P-value* | * |
| **[SERUM]** | **Control** | 2.82 | 1.55 | | | 1.03 | 0.81 | | |
| | **Citalopram** | 1.5 | 1.34 | 0.0001 | *** | 0.58 | 0.6 | 0.0061 | ** |
| | **Oxybutynin** | 1.57 | 1.48 | 0.0003 | *** | 0.67 | 0.61 | 0.0845 | ns |
| | **Metoprolol** | 1.48 | 1.26 | 0.0003 | *** | 0.58 | 0.56 | 0.0118 | * |
| | **Oxycodone** | 1.42 | 1.35 | <0.0001 | **** | 0.56 | 0.53 | 0.0048 | ** |
| | **High DBI** | 2.01 | 1.74 | 0.0317 | * | 0.82 | 0.93 | 0.3371 | ns |
| **[FIRST PASS]** | **Control** | 2.61 | 1.54 | | | 0.99 | 0.79 | | |
| | **Citalopram** | 0.75 | 0.83 | <0.0001 | **** | 0.31 | 0.37 | <0.0001 | **** |
| | **Oxybutynin** | 1.09 | 0.1 | <0.0001 | **** | 0.48 | 0.43 | 0.0005 | *** |
| | **Metoprolol** | 1.42 | 1.18 | <0.0001 | **** | 0.63 | 0.62 | 0.0192 | * |
| | **Oxycodone** | 1.44 | 1.27 | <0.0001 | **** | 0.63 | 0.59 | 0.0143 | * |
| | **High DBI** | 0.97 | 1.1 | <0.0001 | **** | 0.42 | 0.43 | <0.0001 | **** |

SUPPLEMENTARY FIGURES

*Figure S1: Single distribution of fenestration diameters presented in Figure 4 (serum concentrations)*

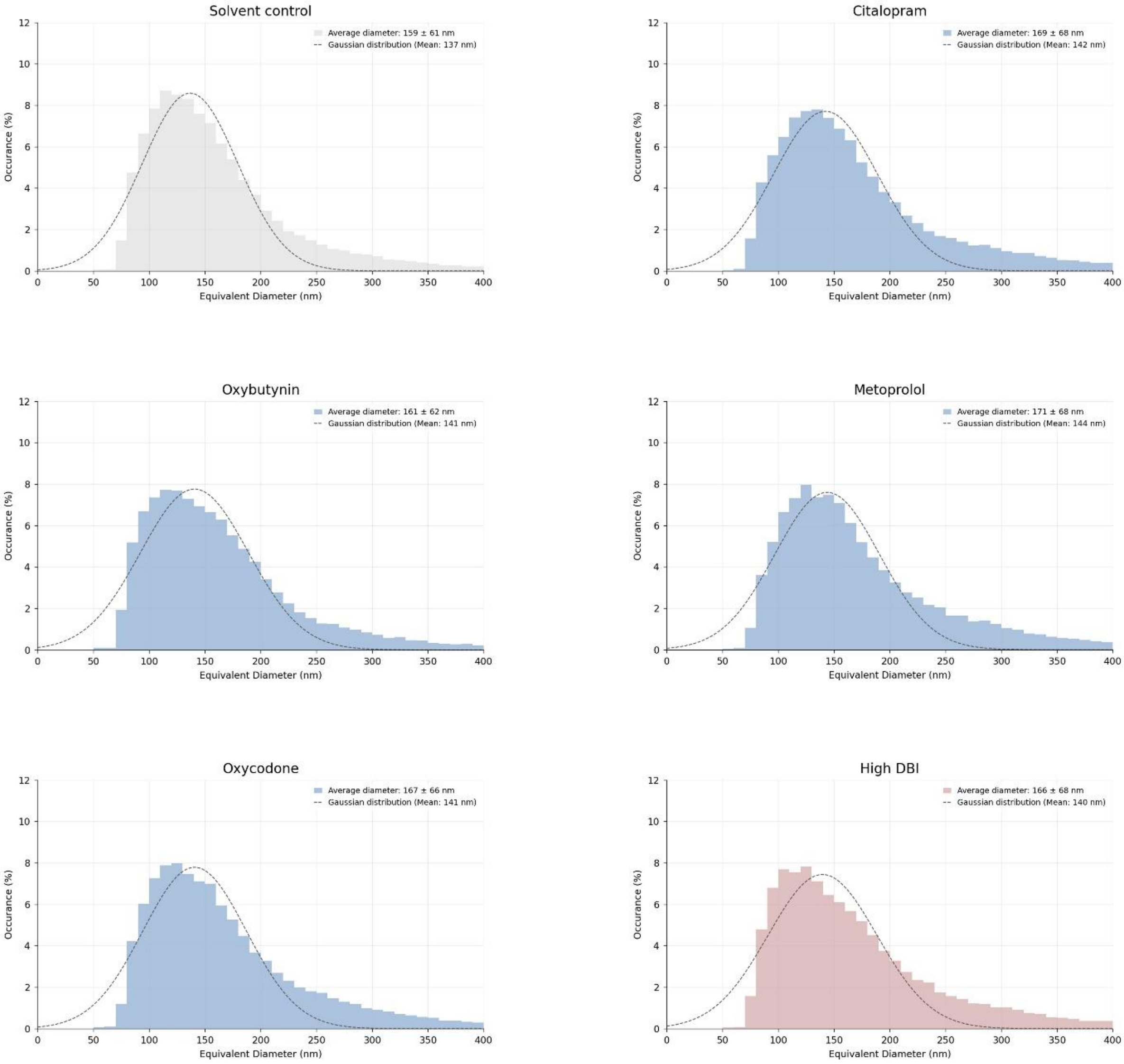

*Figure S2: Single distribution of fenestration diameter presented in Figure 4 (first-pass concentrations)*

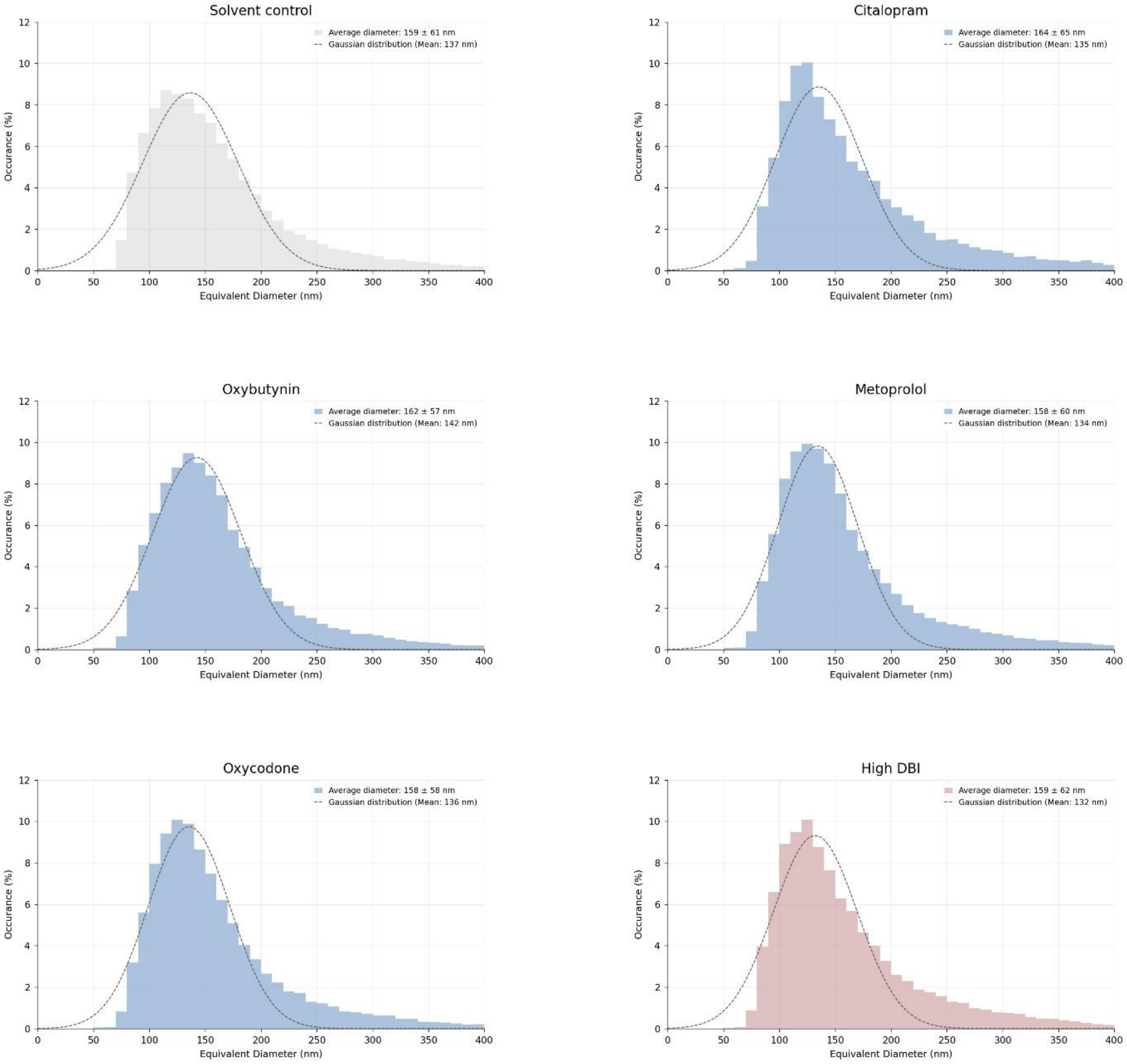
